# TNF signalling fine-tunes Langerhans cell transcriptional programmes mediating adaptive immunity

**DOI:** 10.1101/2021.02.07.430111

**Authors:** James Davies, Andres F. Vallejo, Sofia Sirvent, Gemma Porter, Kalum Clayton, Yamkela Qumbelo, Patrick Stumpf, Jonathan West, Clive M. Gray, Nyaradzo T.L Chigorimbo-Murefu, Ben MacArthur, Marta E Polak

## Abstract

Langerhans cells (LCs) reside in the epidermis as a dense network of immune system sentinels, coordinating both immunogenic and tolerogenic immune responses. To determine molecular switches directing induction of LC immune activation, we performed mathematical modelling of gene regulatory networks identified by single cell RNA sequencing of LCs exposed to TNF, a key pro-inflammatory signal produced by the skin. Our approach delineated three programmes of LC phenotypic activation (immunogenic, tolerogenic or ambivalent), and confirmed that TNF enhanced LC immunogenic programming. Through regulon analysis followed by mutual information modelling, we identified *IRF1* as the key transcription factor for the regulation of immunogenicity in LCs. Application of a mathematical toggle switch model, coupling *IRF1* with tolerance-inducing transcription factors, determined the key set of transcription factors regulating the switch between tolerance and immunogenicity, and correctly predicted LC behaviour in LCs derived from different body sites. Our findings provide a mechanistic explanation of how combinatorial interactions between different transcription factors can coordinate specific transcriptional programmes in human LCs, interpreting the microenvironmental context of the local tissue microenvironments.

## INTRODUCTION

Langerhans cells (LCs) act as immune sentinels at the epidermis and, through antigen presenting function, are responsible for maintaining tissue immune homeostasis (Nestle *et al*., 2009). In the steady-state, a network of LCs resides within the dense assembly of epidermal keratinocytes (KCs), sensing the environment and capturing antigens through intercellular extension and retraction of dendritic processes (Clausen and Stoitzner, 2015). On encounter with antigen, LCs cease to phagocytose and instead upregulate pathways associated with maturation, including MHC II antigen presentation, T cell co-stimulation and migration to local lymph nodes for priming of T cell immunity (Reis e Sousa, Stahl and Austyn, 1993). In the context of diverse signalling from the external environment and epidermal microenvironment, LCs can promote immunogenic responses to protect against harmful pathogens, or promote tolerogenic responses to prevent unwarranted inflammation to self-antigen and innocuous agents (Polak *et al*., 2017)(Sirvent *et al*., 2020)(Clayton *et al*., 2017)(Banchereau and Steinman, 1998). The correct orchestration of immunogenic vs tolerogenic responses by LCs to the different stimuli they encounter is therefore expected to be fundamental to the maintenance of skin health. However, the molecular mechanisms for this decision-making process are largely unknown.

Recent investigations by us and others characterised plasticity in LC-driven adaptive immune responses, dependent on LC activation state and signalling from the skin microenvironment. In the absence of inflammation, migratory LC are marked with enhanced expression of immunocompetency genes and they preferentially promote induction of Th2 CD4+ T cell responses (Sirvent *et al*., 2020)(Polak *et al*., 2014)(Polak *et al*., 2012)(Klechevsky *et al*., 2008), and tolerogenic FOXP3+ Treg responses (Davies *et al*., 2019)(Seneschal *et al*., 2012)(Kitashima *et al*., 2018). In contrast, with TNF signalling, LC immunogenicity is enhanced (Barker *et al*., 1991). TNF is a skin proinflammatory cytokine, which is produced by epidermal KCs, as well as dermal DCs, plasmacytoid DCs (pDCs) and NK cells (Cumberbatch, Dearman and Kimber, 1997)(Singh *et al*., 2016)(Hjorton *et al*., 2018) in response to immunogenic stimuli. TNF stimulation of migratory LC heightens their ability to drive CD8 T cell activity through antigen cross-presentation (Sirvent *et al*., 2020)(Polak *et al*., 2014)(Polak *et al*., 2012). Consistent with enhanced T cell activation, TNF stimulation promotes the upregulation of costimulatory molecules and maturation markers in LC, as well as promoting migration (Berthier-Vergnes *et al*., 2005)(Cumberbatch *et al*., 1999)(Epaulard *et al*., 2014). Furthermore, TNF signalling augments LC mediated anti-viral immunity to human immunodeficiency virus (HIV), Influenza and Epstein-Barr virus (EBV) antigen (Epaulard *et al*., 2014)(Polak *et al*., 2017).

Immune cell function and changes in behaviour, such as the ones observed for LCs, are encoded by unique transcriptomic expression profiles – transcriptional programmes (Sirvent *et al*., 2020)(Xue *et al*., 2014)(Werner, Barken and Hoffmann, 2005)(Hoffmann *et al*., 2002). These transcriptional programmes are coordinated by gene regulatory networks (GRNs) in which transcription factors (TFs) cooperate to define a specific, signal-induced immune outcome (Singh, Khan and Dinner, 2014)(Lin *et al*., 2015). Importantly, interactions with the external environment, tissue status (health or disease) or local microenvironmental signalling, can directly regulate the behaviour of GRN, alter transcriptional programmes and induce functional changes in cells.

Thus, we hypothesised that the decision-making process of LC-driven immunity is determined by the context of the signalling environment, through alteration of transcriptional programmes underpinning LC activation. We assumed that, while spontaneous migration in the absence of pro-inflammatory signalling reflects the scenario in which LCs mediate peripheral immune homeostasis, TNF signalling favours immunogenicity. We sought to identify specific TFs defining immunogenic and tolerogenic programmes in LCs and to determine the regulatory interactions between the phenotype-defining TFs. Combining single cell transcriptome analyses with a published toggle switch ordinary differential equation (ODE) model defining two divergent sets of TF expression, containing self-amplification and mutual inhibition (Huang *et al*., 2007), we identified regulatory modules defining immunogenic (*IRF1, IRF4*) and tolerogenic (*IRF4, RELB, ELK1, KRAS, SOX4*) LC phenotypes. The model was used to predict LC transcriptional programmes across abdominal skin, breast skin and foreskin-derived migrated LC, and provides a mechanistic explanation of how combinatorial interactions between different transcription factors can coordinate tissue and activation-specific transcriptional programmes in human LCs.

## RESULTS

### TNF enhances immunogenic transcriptional programming in migratory LC

In order to mediate protective immune responses, epidermal LCs must respond appropriately to environmental cues. To investigate transcriptional programmes induced by epidermal pro-inflammatory cytokines in LCs, we performed single cell analysis of human primary migrated LCs exposed to 24h stimulation with TNF vs unstimulated control. Clustering and dimensionality reduction analysis of 737 cells (UMAP, ScanPy, version=1.5.0) revealed that LC migrated from abdominal skin and cultured in the presence or absence of TNF contained a predominant large cluster, confirmed to be LCs through high expression of MHC II genes (CD74, *HLA-DRB1, HLA-DRB5*), as well as two additional populations identified to be melanocytes (TYRP1, TYR) and T cells (CD3D) (Logistic regression, ScanPy pipeline, version=1.5.0), which were removed from downstream analysis (Supplementary figure 1A-C). The heterogeneity of the 737 migrated LCs cultured with or without TNF (unstimulated = 375, TNF stimulated = 362) was then analysed. Overall, the cells appeared relatively homogeneous, consisting of one overall large population of LCs comprising sub clusters of unstimulated and TNF stimulated LCs, which appear to diverge away from each other (Figure 1A). Differentially expressed gene (DEG) analysis comparing migrated LCs with and without TNF identified 61 genes upregulated in unstimulated LCs and 87 genes upregulated in TNF stimulated LCs (MAST, adj.p-value<0.05, Supplementary figure 1D). Gene ontology analysis of the 61 genes upregulated in unstimulated LCs showed they were associated with secretion by cell (adj. P-Value=5.3E-3) and regulation of the immune response (adj. P-Value=5.3E-3, Figure 1B&C), with the latter influenced by the upregulation of the TF *KRAS* (Figure 1B, Supplementary figure 1E). Gene ontology analysis for the 87 genes upregulated in migrated TNF-stimulated LCs revealed association with cytokine-mediated signalling pathways (adj. P-Value=2.2E-7) and positive regulation of alpha-beta T cell activation (adj. P-Value=1.5E-4)(Figure 1B&C). This signature was influenced by the expression of the TF *IRF1*. Using an immunogenic gene signature comprising genes upregulated in TNF stimulated LC (0hr-24hr DEGs) from bulk RNA-seq data (Sirvent *et al*., 2020) and a previously described tolerogenic gene signature comprising genes associated with dendritic cell tolerogenic function (Davies *et al*., 2019) (Supplementary Table 1), z-scores representing the activation of each programme were calculated for individual LCs (Figure 1D). Differences in z-score (Immunogenic-tolerogenic) were calculated, with the positive values reflecting LCs with increased immunogenic signature expression. This revealed that overall, TNF stimulated LCs display an enhancement for the immunogenic signature (Median = 0.02107) compared to unstimulated (Median = -0.09535, unpaired t-test, p=<0.001)(Figure 1E). Overall TNF stimulation enhances LC transcriptomic programmes associated with immunogenic responses and T cell activation.

**Figure 1.**
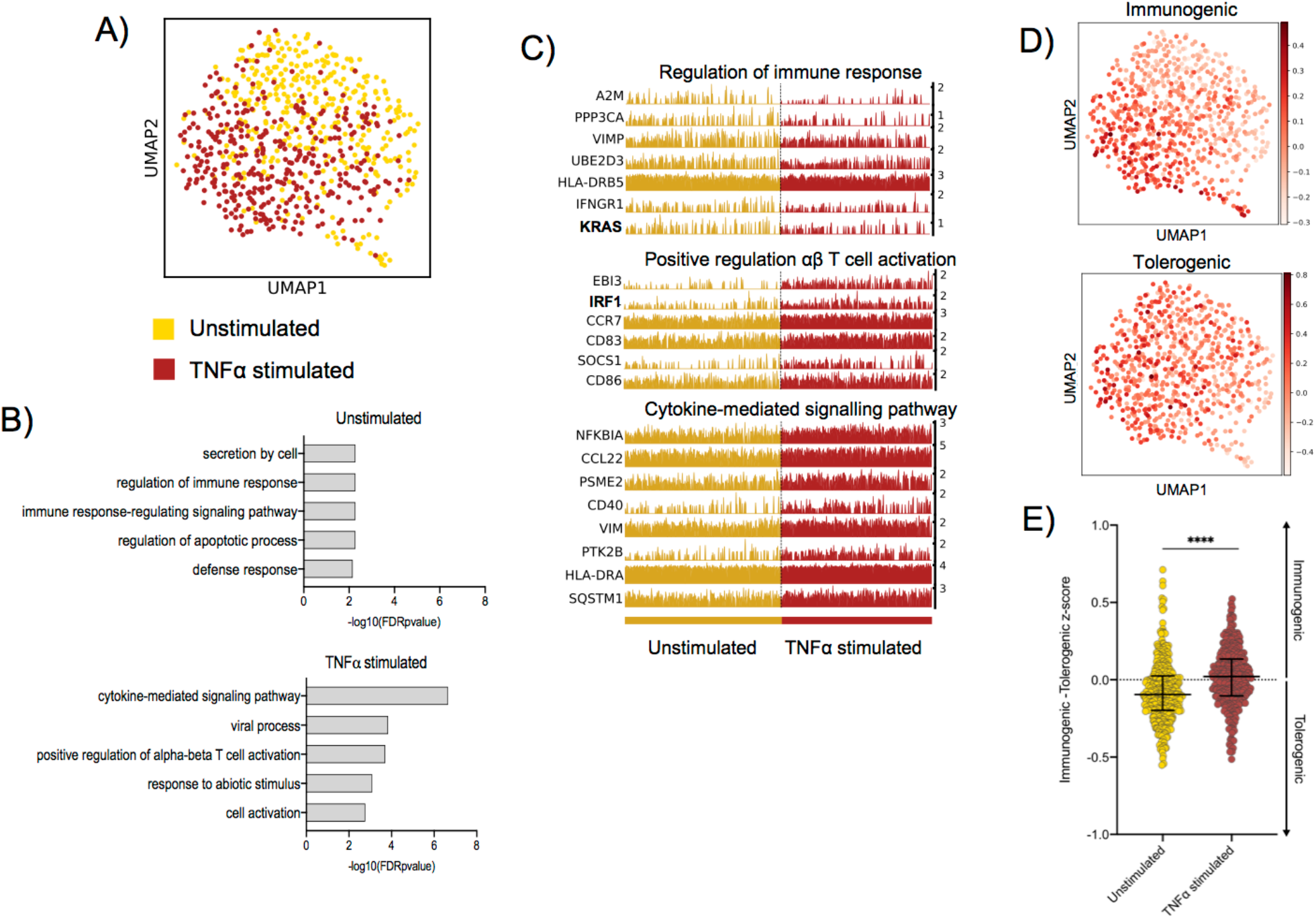
TNF enhances immunogenic transcriptional programming in migratory LC. **A**. UMAP dimensionality reduction analysis of scran-normalised single cell data from unstimulated (375) and TNF stimulated (362) migrated LCs originating from the same donor. **B**. Gene ontology analysis (Toppgene) for the 61 upregulated DEGs in unstimulated migrated LCs and 87 upregulated DEGs in TNF stimulated migrated LCs (FDR corrected p=<0.05) **C**.Trackplots displaying genes included in ontologies upregulated in unstimulated migrated LC (regulation of immune response) and TNF stimulated migrated LC (positive regulation of αβ T cell activation and cytokine-mediated signalling pathway). **D** UMAP marker plots displaying immunogenic z-scores and tolerogenic signature z-scores in individual LC. Immunogenic z-scores were derived from the expression of genes identified to be upregulated in TNF stimulated LC (0hr-24hr DEGs) from bulk RNA-seq data (Sirvent *et al*., 2020). Tolerogenic signature z-scores were derived from the expression of genes associated with dendritic cell tolerogenic function (Davies *et al*., 2019). **E**.Differences in z-scores (immunogenic – tolerogenic z-score) were calculated to compare the proportion of unstimulated and TNF stimulated migrated LC displaying an elevated immunogenic profile. Unpaired t-test, ****=p<0.001.

### *IRF1* expression controls immunogenic transcriptional programming

To identify the key TF regulators of programming in unstimulated vs TNF-stimulated migrated LC, SCENIC (Aibar *et al*., 2017) single cell regulatory network inference analysis was performed (Figure 2A, z-score enrichment ≥0.1). Here, TNF stimulated LC displayed enrichment of the *IRF1* regulon (Figure 2B, z-score=0.2), which, along with the upregulated expression of *IRF1* from DEG analysis (Figure 2C, MAST), strongly highlighted this TF as being a candidate critical for immunogenic LC programming. In unstimulated LC, the most enriched regulon was *SOX4*, although this enrichment was more moderate (Figure 2A, z-score=0.1). Interestingly, *IRF4*, which has been demonstrated to be critical for both LC immunocompetent and tolerogenic programming (Sirvent *et al*., 2020)(Davies *et al*., 2019), displayed homogenous regulon enhancement and expression across both populations (Supplementary Figure 2A-B).

**Figure 2.**
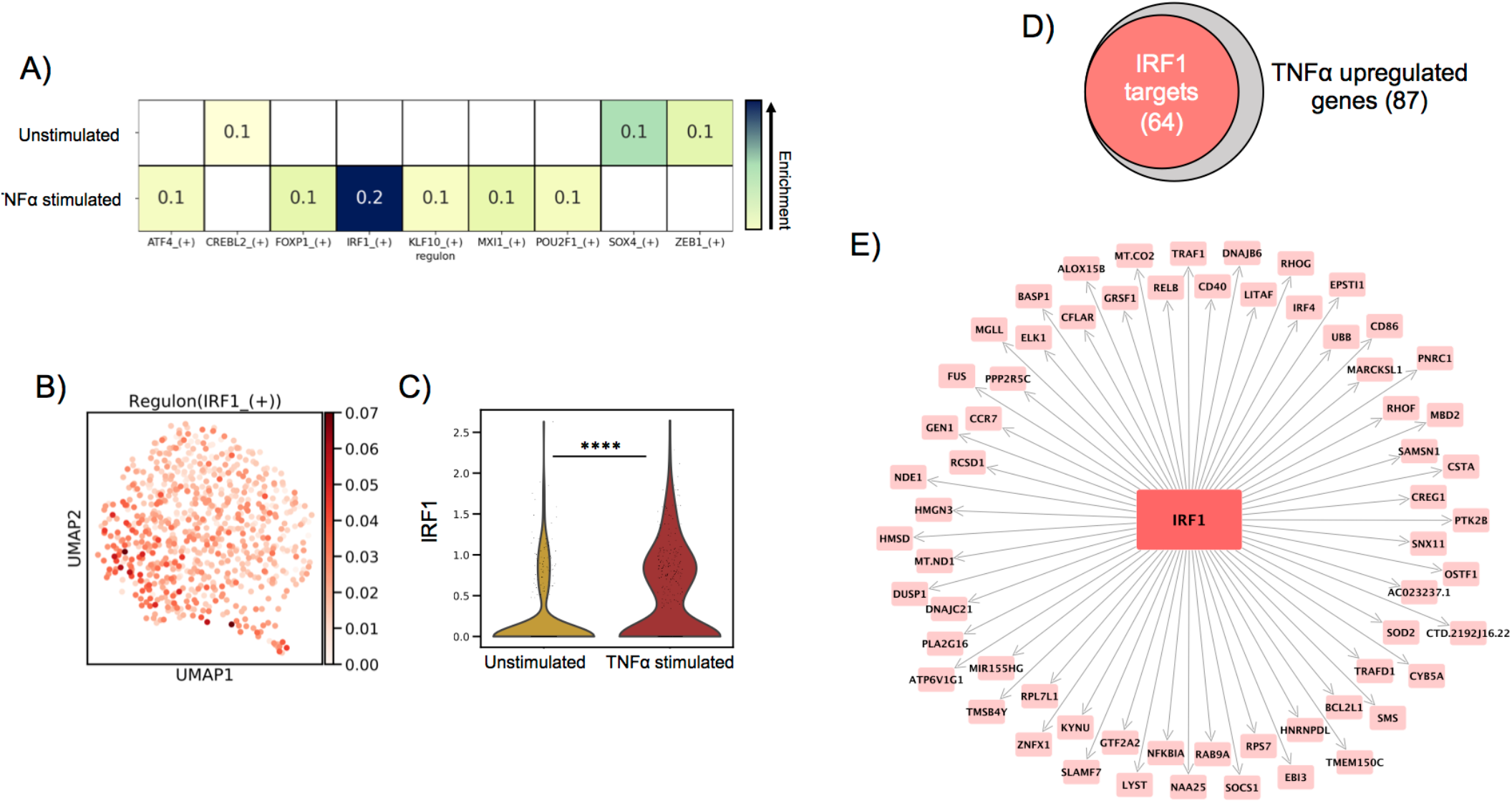
IRF1 expression controls immunogenic transcriptional programming. **A**. SCENIC regulatory network and inference clustering analysis revealed TF regulons which were enriched in unstimulated and TNF stimulated migrated LCs. Z-score heatmap (yellow-> blue) of enriched regulons are displayed (z-score>0.1). **B**. UMAP marker plot displaying *IRF1* regulon enrichment (z-score) in individual LCs. **C**. Violin plot displaying the level of transcriptomic expression of *IRF1* in unstimulated and TNF stimulated migrated LCs, MAST, ****=p<0.001. **D** Venn Diagram displaying the overlap in TNF stimulated LC upregulated genes identified to be targets of *IRF1* in PIDC analysis (edge weight >1). **E** PIDC network graph displaying IRF1 targets (edge weight >1) identified within TNF stimulated LC upregulated genes.

Whilst unstimulated LCs displayed significant upregulation of *KRAS* and enrichment of the *SOX4* regulon, these findings were relatively weak and less exclusive to unstimulated LC in contrast to the clear upregulation of *IRF1* in TNFα stimulated LC (Figure 2B&C, Supplementary Figure 2B). We therefore explored whether these TFs acted in accordance with core TF mediators of programming in migrated LC which have previously been associated with coordinating immunocompetent and tolerogenic regulation, including *IRF4, RELB, ELK1, KLF6* and *HMGN3* (Davies *et al*., 2019). Using partial information decomposition analysis (PIDC) (Chan, Stumpf and Babtie, 2017) gene regulatory network inference of the 61 genes upregulated in unstimulated migrated LC, along with *KRAS, SOX4, IRF4, RELB, ELK1, KLF6* and *HMGN3*, a directed PIDC (TF -> target gene edges only) network graph depicting regulatory interactions between TFs and target genes was generated (correlation score >1.5). Here, *KRAS* and *SOX4* could be observed to be components of a highly interconnected regulatory hub with *IRF4, RELB* and *ELK1* (Supplementary Figure 2C). This regulatory hub could be associated with controlling the expression of 33 unstimulated LC upregulated genes, highlighting its importance for transcriptional programming in unstimulated LC (Supplementary Figure 2D). PIDC analysis was also performed to identify targets of *IRF1* within the TNF-upregulated gene list to discern the TFs influence on the transcriptomic programming on TNF-stimulated LC. Here, 64/87 (74%) TNF-upregulated genes were identified to be targets of *IRF1* (Figure 2D-E). Furthermore, PIDC analysis of *IRF1* along with the core migrated LC TFs and the 87 genes upregulated in TNF stimulated LC, suggested *IRF1* upregulation added an additional layer of regulation beneath the core network of *ELK1, RELB, IRF4* and *HMGN3* to mediate immunostimulatory programming. (Supplementary figure 2E).

### A toggle switch mathematical model predicts immunostimulatory vs tolerogenic

#### LC phenotypes from single cell transcriptomic data

Single cell analysis revealed distinct programming of unstimulated and TNF-stimulated LCs determined by differentially regulated TFs. To explore how the balance of LC phenotypes is controlled, we utilised a tri-stable toggle switch ODE model in which different activation programmes can be described based on the expression of a selected number of programme (immunogenic vs tolerogenic) defining TFs (Huang *et al*., 2007). The ODE model contains 2 equations which each represent the activation of immunogenic (*I)* and tolerogenic (*T*) programmes, respectively (Figure 3A). Each equation contains 3 terms, which represent auto-amplification (dotted box), cross-inhibition of opposing programmes (dashed box) and first order state decay (solid box). The model therefore assumes that the regulatory programmes that define each programme auto-amplify their own expression, whilst inhibiting the expression of the opposite programme. The tri-stable model describes a phenotypic ‘attractor landscape’ in which LCs can fall into an immunogenic (A), a tolerogenic (B) or an ambivalent (C, equal ability to stimulate tolerogenic and immunogenic responses) state (Figure 3B). In the phase portrait, (A) and (B) therefore represent states in which the expression of TFs from either programme is dominant over the other, whilst (C) represents a state in which there is balanced expression of both immunogenic and tolerogenic programmes. The model can therefore be utilised on single cell data to predict the phenotypic state of individual LCs by plotting single LC trajectories in state space using single cell expression data z-scores of phenotype-defining TFs.

**Figure 3.**
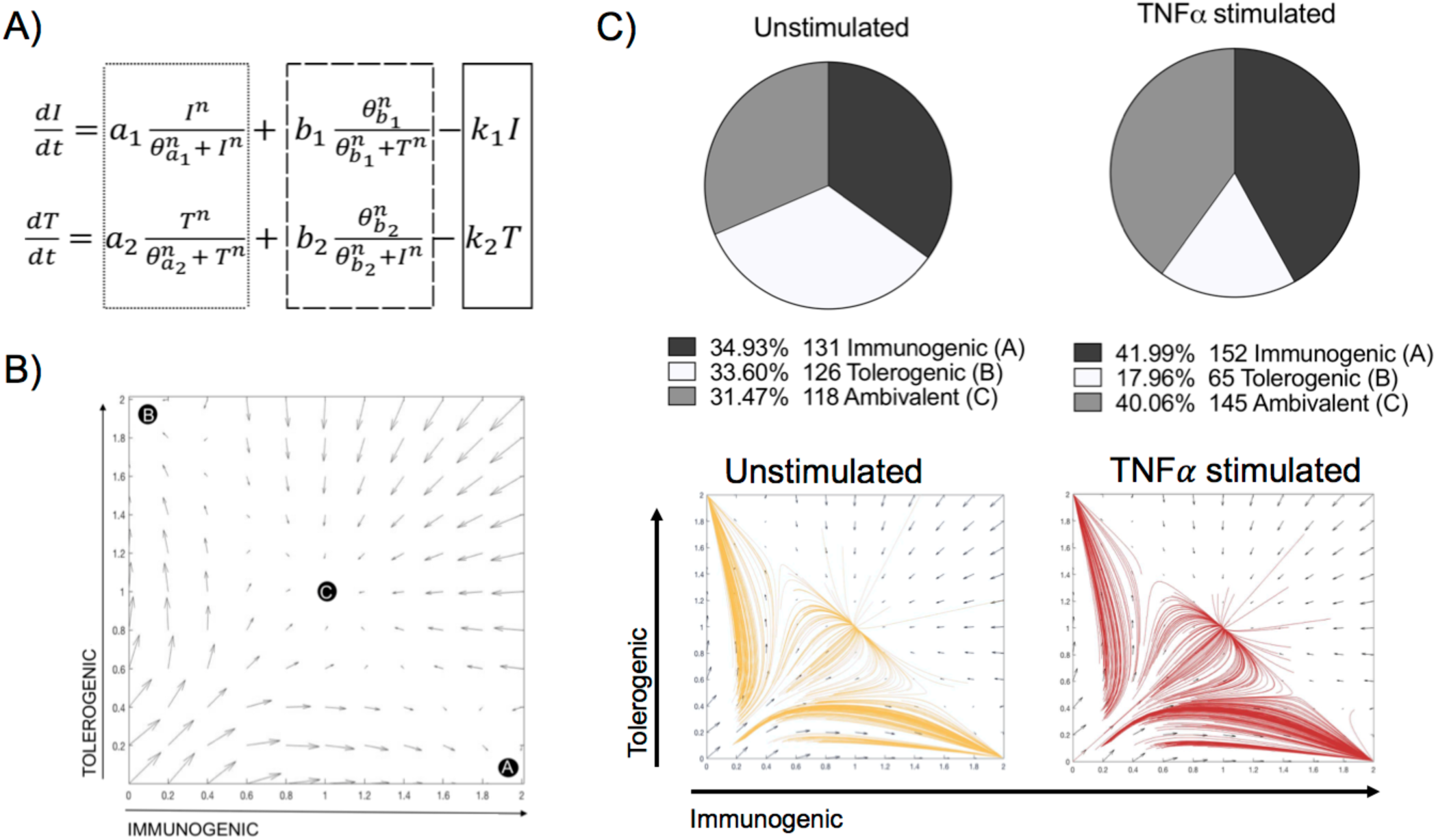
A toggle switch mathematical model predicts immunogenic vs tolerogenic LC phenotypes from single cell transcriptomic data. **A**. Dynamical system representing the activation of immunogenic (I) and tolerogenic (T) programmes in LCs. The dotted box represents terms describing the auto-amplification of each respective programme. The dashed box represents terms describing the cross-inhibition from opposing programmes, whilst the solid box depicts the first-order decay rate (k) for each programme. **B**. Phase portrait of the toggle switch model in which the two programmes (immunogenic and tolerogenic) auto-amplify their own expression and are mutually repressive. Black circles (A, B and C) represent end points for trajectories at stable attractors representing an immunogenic programme (A), tolerogenic programme (B) or an ambivalent programme (C). **C**.Pie charts summarising the numbers and percentages of LC assigned to each phenotype through utilising the toggle-switch model for trajectory plotting. For each trajectory, representing an individual unstimulated or TNF-stimulated LC, the x-axis represents z-scores combining normalised *IRF1/IRF4* expression values; the y-axis represents the z-scores combining *SOX4, KRAS, IRF4, RELB* and *ELK1* expression values. Z-scores were scaled to fit phase portrait boundaries.

The model has been systematically tested by iterative application of distinct transcription factor combinations (Supplementary Table 2). For defining the immunogenic phenotype, *IRF1* alone or in combination with *IRF4* was tested. The inclusion of *IRF4* for immunogenic regulation was based on previous analysis demonstrating the importance of *IRF4* for both immunizing and tolerizing T cell activation, as well as immunocompetent LC programming (Vander Lugt *et al*., 2017)(Sirvent *et al*., 2020), which was supported by our PIDC analysis which revealed extensive interconnectivity of these TFs. For defining the tolerogenic phenotype, combinations of *KRAS, SOX4, IRF4, RELB* and *ELK1* were investigated. To define the best model however, we reflected on which models best followed the hypothesis that unstimulated migrated LCs are mutually efficient at inducing immunogenic and tolerogenic responses. Likewise, the model predictions would need to reflect the differences in z-score signature enrichment observations, in which TNF stimulated LC exhibited an increase in immunogenic signatures and a reduction in tolerogenic signatures (Figure 1E). Overall, many model iterations depicted the observations that the TNF-stimulated LC population contain increased quantities of immunogenic LCs (Supplementary Table 2). However, model 14, in which both *IRF1* and *IRF4* depicted the immunogenic phenotype and *KRAS, SOX4, IRF4, RELB* and *ELK1* depicted the tolerogenic phenotype, was best at predicting results in line with both criteria (Figure 3C, Supplementary Table 2). Here, the relative quantities of immunogenic (34.93%), tolerogenic (33.60%) and ambivalent (31.47%) LCs in unstimulated LCs was equal, whilst TNF stimulated LCs displayed an increase in immunogenic (41.99%) and ambivalent (40.05%) programmed LCs and a decrease in tolerogenic (17.96%) LCs.

### IRF1/IRF4 toggle-switch determines body-site specific differences in LC

#### immunogenic programming

We next sought to validate the power of the model to predict differences in transcriptomes from LCs of independent datasets, including a single cell dataset of migrated breast skin-derived and foreskin-derived LC. Comparative analysis of z-score enrichment for immunogenic vs tolerogenic signatures revealed that foreskin LCs more frequently display a predominant immunogenic phenotype (Figure 4A, p=0.039). This enhanced immunogenic programme in foreskin LCs could be seen in the expression of inflammatory pathway-associated transcripts, which importantly, included *IRF1* (Figure 4B). The model was then applied, using the same parameters and TFs as in Figure 3C, to test model predictions of immunogenic, tolerogenic and ambivalent populations amongst breast skin and foreskin-derived LC. Here the model predicted breast skin LCs to be 9.35% immunogenic, 37.92% tolerogenic and 52.73% ambivalent, whilst foreskin LCs were predicted to be 16.67% immunogenic, 29.17% tolerogenic and 54.17% ambivalent (Figure 4C). Model predictions of increased immunogenicity in foreskin LC therefore reflected transcriptomic observations in which foreskin derived LC display enhancement of immunogenic programming.

**Figure 4.**
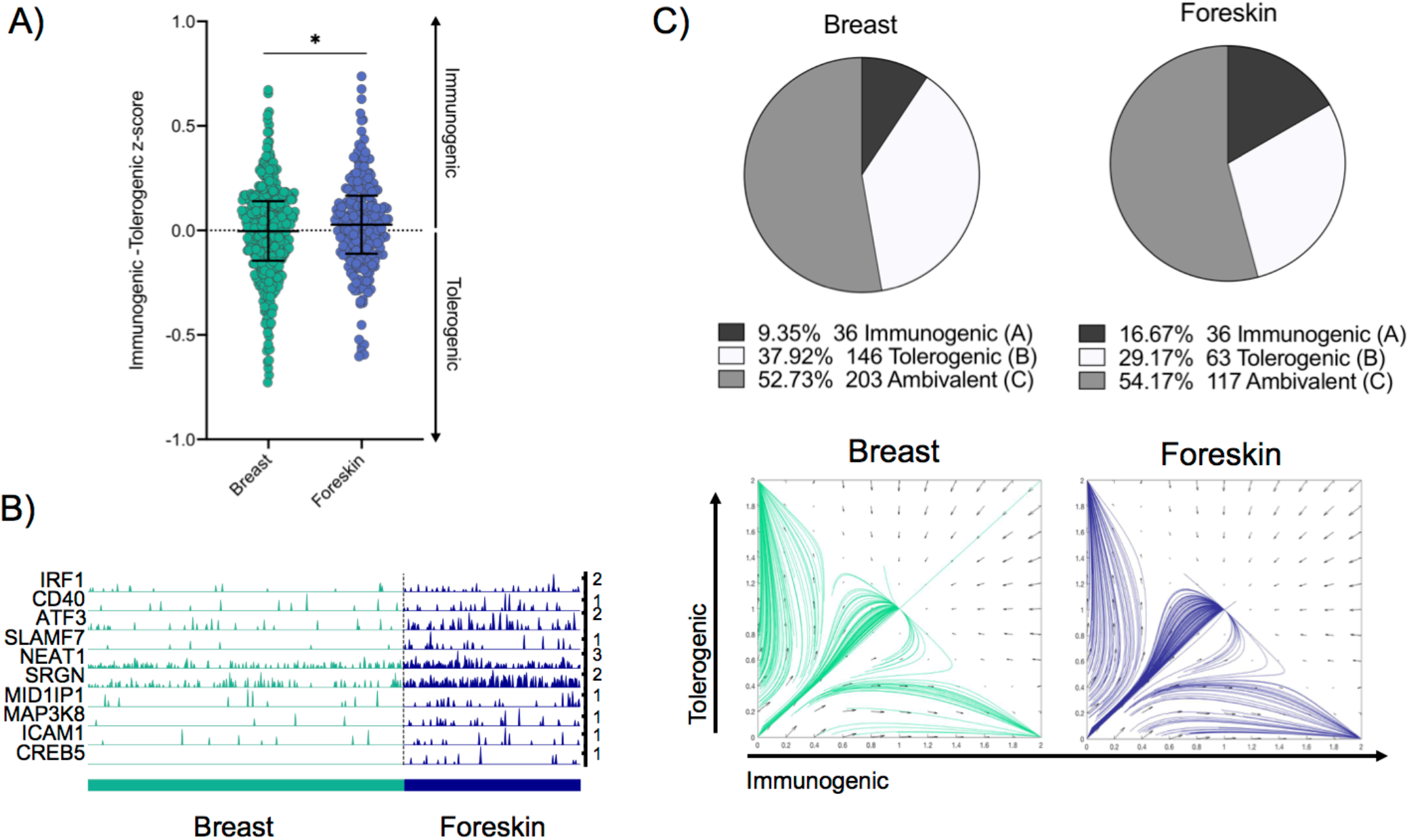
IRF1/IRF4 toggle-switch determines body-site specific differences in LC immunogenic programming. **A**. Differences in z-scores (immunogenic – tolerogenic z-score) were calculated to quantify the proportion of breast derived skin and foreskin migrated LCs that display elevated immunogenic profiles. Unpaired t-test, *=p<0.05. **B**. Trackplots comparing the expression of transcripts associated with immunogenic LC function across breast skin derived and foreskin migrated LCs. **C**. Pie charts summarising the numbers and percentages of LC assigned to each phenotype through utilising the toggle-switch model for trajectory plotting. For each trajectory, representing an individual breast-derived or foreskin-derived migratory LC, the x-axis represents z-scores combining normalised IRF1/IRF4 expression values; the y-axis represents the z-scores combining SOX4, KRAS, IRF4, RELB and ELK1 expression values. Z-scores were scaled to fit phase portrait boundaries.

## DISCUSSION

Immune cell function and behaviour are encoded by unique transcriptomic expression profiles – transcriptional programmes (Xue *et al*., 2014). Changes in the transcriptional programmes, which reflect status of health or disease or environmental signalling, are coordinated by gene regulatory networks (GRNs) in which transcription factors (TFs) play an essential role (Singh, Khan and Dinner, 2014)(Lin *et al*., 2015). However, large scale investigations into the activity of individual GRN components and interactions between specific modules which underlie different transcriptomic programmes, and in particular the kinetics in which those programmes are executed, are difficult to investigate using functional *in vitro* methods (Ay and Arnosti, 2011). Therefore, mathematical modelling techniques are increasingly being utilised to counter this problem and include methods such as ordinary differential equation (ODE) modelling and Petri net modelling (Loriaux and Hoffmann, 2012)(Livigni *et al*., 2018). Mathematical modelling can permit investigations of dynamic biological systems *in silico* to assess how different molecular signals can alter regulatory network behaviour. For example, Petri net modelling has revealed the LC IRF-GRN underlying immunogenic immune activation in response to different stimuli (Polak *et al*., 2017). However, Signalling Petri Net (SPN) and similar methods allow only qualitative assessment of network behaviour, and limit the strength of predictions. In contrast, ODE modelling has allowed exploration of small TF networks and specific network elements, such as positive feedback based switches, which can define cell lineage determination and operon activation (Huang *et al*., 2007)(Gardner, Cantor and Collins, 2000).

In GRNs, TFs act in concert with each other to coordinate different expression programmes. However, specific cellular phenotypes are determined by the increased expression of specific phenotype-defining TFs. For example, in macrophages, whilst *NFKB1, JUNB* and *CREB1* define core programmes of activation, *STAT4* is specifically upregulated in the context of chronic inflammation, which correlates with increased expression of a unique gene expression programme (Xue *et al*., 2014).

The plasticity of migrated LC to induce both immunogenic and tolerogenic adaptive T cell responses (Sirvent *et al*., 2020)(Polak *et al*., 2014)(Polak *et al*., 2012)(Klechevsky *et al*., 2008)(Davies *et al*., 2019) has revealed the complexity in discerning the decision-making process of LCs to drive either immunogenic or tolerogenic responses and has highlighted the question as to how LCs skew T cell activation to favour responses that are preferential in different biological contexts such as inflammation. Here we analysed single cell transcriptomic data arising from unstimulated and TNF-stimulated migrated LC, to discern the divergent programming of LCs in response to inflammatory stimuli and uncover critical TFs which govern immunogenic gene regulation.

The epidermal inflammatory cytokine TNF is a powerful mediator of inflammation and its effects on enhancing LC activation and programming of immunogenic T cells has previously been demonstrated (Sirvent *et al*., 2020)(Stoitzner *et al*., 1999),(Théry and Amigorena, 2001). Here we demonstrate at the single cell level that, compared to unstimulated LCs, TNF causes divergent transcriptional programming characterised by upregulation of genes associated with inflammatory cytokine signalling processes and T cell activation, thus reflecting their enhanced immunogenic function *in vitro*. Interestingly whilst the effects of TNF were clear, there was still significant overlap between the stimulated and unstimulated populations, suggesting that common transcriptomic features, likely associated with migration and immunocompetency, were still present. Importantly, as highlighted from our analysis, the TF *IRF1* was revealed to be a critical component of the TNF-enhanced transcriptomic programme, which appeared to be projected onto the core migrated LC transcriptional network to enhance immunogenic programming. The association of *IRF1* with inflammatory pathway activation has been observed in other systems. In DCs, TLR-9-mediated IRF1 induction leads to the induction of IFN*γ* and interferon-stimulated genes, driving efficient anti-viral immune responses (Schmitz et al., 2007). *IRF1* activation in macrophages is associated with the polarisation of macrophages towards the pro-inflammatory M1 phenotype (Chistiakov et al., 2018). In fibroblast like synoviocytes (FLS), which are implicated in the inflammation in rheumatoid arthritis, TNF-mediated induction of *IRF1* leads to induction of inflammatory mediators, such as IFN*γ* (Bonelli et al., 2019). In contrast, *IRF4* has been conclusively demonstrated as a transcription factor critical for LC immunocompetent programming and DC capacity to induce immunogenic T cell activation (Vander Lugt *et al*., 2017)(Sirvent *et al*., 2020). Therefore, we hypothesised that together, *IRF1* and *IRF4* complementarily coordinate LC immunogenic programming. In our model, while expression of IRF4 induces LC maturation and immune-competence, expression of IRF1, induced by TNF signalling, fine-tunes the programmes towards immunogenicity. Additionally, we revealed that in unstimulated LCs, *KRAS* and *SOX4* interact with components of a core network of TFs enhanced upon LC migration (*IRF4, RELB* and *ELK1*), previously demonstrated to be responsible for immunocompetent and tolerogenic regulation (Davies *et al*., 2019). This revealed the preference by unstimulated, migrated LC, for homeostatic regulation as compared to the immunogenic regulation enhanced in TNF-stimulated, migrated LC.

*In vivo* analysis of LC behaviour in humans is unfeasible and *in vitro* methods to observe phenotypic behaviours are constrained. The utilisation of mathematical modelling is therefore fundamental to augmenting comprehension of phenotypic programmes of LC *in situ*. Importantly, interpretation of transcriptomic observations in light of a well-established toggle-switch model of general cell fate specification (Huang *et al*., 2007) permitted an unprecedented opportunity to explore the determinants regulating immunogenic vs tolerogenic programmes in LC. Analysis of this model indicates that 3 stable phenotypes are possible, which could reflect the phenotypic landscape in which LC can adopt predominantly immunogenic or tolerogenic programmes, or an intermediate ambivalent programme, in which immunogenic and tolerogenic activation are mutually present and in balance. Such “multilineage priming” is common in cell fate switches and may have an important role in regulating LC fate decisions. Using the model in which *IRF1/IRF4* determine immunogenicity and *KRAS/SOX4/IRF4/RELB/ELK1* determine tolerogenicity, we demonstrated that model predictions were reflective of our transcriptomic data. Moreover, the model allowed prediction of *in vitro* phenotypic features of enhanced immunogenicity in TNF stimulated LC (Sirvent *et al*., 2020)(Polak *et al*., 2014).

The foreskin microenvironment is associated with increased need for effective anti-microbial responses and is reported to be a pro-inflammatory/ immunologically active tissue marked by elevated pro-inflammatory cytokines and infiltration of effector immune cells (Prodger *et al*., 2012)(Sennepin *et al*., 2017)(Zhou *et al*., 2011)(Gray *et al*., 2020). Apart from baseline and mitogen-induced TNF and IFN*γ* secretion by foreskin CD8 T cells being higher than levels secreted by CD8 T cells in the blood (Prodger *et al*., 2012), the foreskin is most likely in a consistent state of inflammation being driven by infiltrating T cells and elevated LC’s upon exposure to a multitude of microbial stimuli (Gray *et al*., 2020). These inflammatory-associated characteristics of the foreskin site were reflected in transcriptomic observations made during comparison of LC derived from breast skin and foreskin, in which immunogenic programming was enhanced in foreskin LC. Here, we again showed that model predictions were reflective of transcriptomic observations, highlighting the power of the model across anatomically diverse LC datasets.

Overall, we have shown that epidermal signalling, such as pro-inflammatory TNF, can modulate the proportion of LCs exhibiting different immunological programmes. This may therefore, reflect how LCs balance the need for different immunological responses to diverse biological stimuli. Furthermore, we have highlighted specific TF regulators critical for the modulation of both immunogenic and tolerogenic LC programmes, which, when translated into a mathematical model, have demonstrated the potential to predict LC phenotypes across different LC transcriptomic datasets. This opens opportunities to apply the model for predicting LC activation states and behaviour across different biological contexts in health and disease, and provides a tool for assessment of LC activation status in human skin.

## METHODS

### Human LC isolation

Human skin abdominoplasty samples were collected with written consent from donors with approval by the South East Coast -Brighton & Sussex Research Ethics Committee in adherence to Helsinki Guidelines [ethical approvals: REC approval: 16/LO/0999). Fat and lower dermis was cut away and discarded before dispase (2 U/ml, Gibco, UK, 20h, +4°C) digestion. Foreskin tissue was collected from the Medical Male Circumcision HIV prevention programme in Cape Town, South Africa. Tissue was collected with consent and approved by the University of Cape Town [ethics approvals HREC: 566/2012]. Inner and outer foreskin was dissected and processed in an identical manner to the abdominoplasty samples. Migrated LCs were extracted from epidermal explant sheets cultured in media (RPMI, Gibco, UK, 5% FBS, Invitrogen, UK, 100 IU/ml penicillin and 100 mg/ml streptomycin, Sigma, UK) for 48 hours. Migrated LC were purified through fluorescence-activated cell sorting (FACS). TNF stimulated migrated LCs were incubated for 24 hours with 25ng/ml TNFα. Antibodies used for cell staining were pre-titrated and used at optimal concentrations. A FACS Aria flow cytometer (Becton Dickinson, USA) and FlowJo software was used for analysis. For FACS purification LCs were stained for CD207 (anti-CD207 PeVio700), CD1a (anti-CD1a VioBlue) and HLA-DR (anti-HLA-DR Viogreen, Miltenyi Biotech, UK).

### Drop-seq

After FACS purification, single LCs were co-encapsulated with primer coated barcoded Bead SeqB (Chemgenes, USA) within 1 nL droplets (Drop-seq). Drop-seq microfluidic devices according to the design of Macosko *et al*. were fabricated by soft lithography, oxygen plasma bonded to glass slides and functionalised with fluorinated silane (1% (v/v) trichloro(1H,1H,2H,2H-*perfluorooctyl)silane* in HFE-7500 carrier oil). Open instrumentation syringe pumps and microscopes (see dropletkitchen.github.io) were used to generate and observe droplets, using conditions and concentrations according to the Drop-seq protocol, 607 steady-state LC and 208 migrated LC from mastectomy skin were converted into ‘STAMPs’ for PCR library amplification (High Sensitivity DNA Assay, Agilent Bioanalyser) and tagmentation (Nextera XT, Illumina, UK). Sequencing of libraries was executed using NextSeq on a paired end run (1.5×10E5 reads for maximal coverage) at the Wessex Investigational Sciences Hub laboratory, University of Southampton.

### Transcriptomic data analysis

The Drop-seq protocol from the McCarrol lab was followed for converting sequencer output into gene expression data. The bcl2fastq tool from Illumina was used to demultiplex files, remove UMIs from reads and deduce captured transcript reads. Reads were then aligned to human hg19 reference genome using STAR. Analyses was performed using the python-based Scanpy pipeline(version 1.5.0), (Wolf, Angerer and Theis, 2018). High quality barcodes, discriminated from background RNA barcodes, were selected based on the overall UMI distribution using EmptyDrops (Lun *et al*., 2019). Low quality cells, with a high fraction of counts from mitochondrial genes (20% or more) indicating stressed or dying cells were removed. In addition, genes with expression detected in <10 cells were excluded. Datasets were normalised using scran, using rpy2 within python (Lun, Bach and Marioni, 2016). Highly variable genes (top 2000) were selected using distribution criteria: min_mean=0, max_mean=4, min_disp=0.1. A single-cell neighbourhood graph was computed on the first principal components that sufficiently explain the variation in the data using 10 nearest neighbours. Uniform Manifold Approximation and Projection (UMAP) was performed for dimensionality reduction. The Leiden algorithm (Traag, Waltman and van Eck, 2019) was used to identify clusters within cell populations (Leiden r = 0.5, n_pcs=30). Differentially expressed genes (DEGs) between cell clusters were identified using MAST (FDR corrected p-value<0.01, logFC>1). Gene ontology analysis was performed using Toppgene (FDR corrected p-value<0.05), describing biological pathways associated with gene lists. Z-scores for tolerogenic and immunogenic gene signatures were calculated for each single LC. Tolerogenic signature was composed of genes identified to be associated with DC tolerogenic function and previously shown to be enriched in tolerogenic migrated LC (Davies *et al*., 2019). The immunogenic signature was composed of 0-24 hour TNFα stimulated LC upregulated DEGs, identified from bulk RNA-seq data (Sirvent *et al*., 2020). To differentiate LC with predominantly immunogenic or tolerogenic transcriptomic expression profiles immunogenic – tolerogenic z-scores were calculates for each single cell. The more positive the difference in z-score values, the more immunogenic and the more negative the difference in z-scores, the more tolerogenic. Regulatory network inference analysis was performed using single-cell regulatory network inference and clustering (SCENIC) within python (Aibar *et al*., 2017).

### Directional PIDC

Notebooks from Chan et al. were adapted for the analysis and run using Julia V 1.0.5 in Jupyter Notebook. Directional network inference of *IRF1* with TNF stimulated LC upregulated DEGs was performed using PIDC algorithm (Chan, Stumpf and Babtie, 2017) using scran-normalised expression data. Inference of unstimulated and TNF stimulated migrated LC TF -> target networks was performed using scran-normalised expression data of core LC TFs (Davies *et al*., 2019), plus *IRF1* in TNF stimulated LC and the upregulated DEGs for unstimulated and TNF stimulated LCs, respectively. Edge weights were exported, and sorted to include only transcription factors as targets. Hierarchical network was visualised using yED.

### Mathematical modelling

The toggle-switch ODE model was adapted from Huang et al. (Huang *et al*., 2007), in which the observed functional interactions are depicted in an ‘influence’ network, rather than molecular mechanisms of interaction. The model is constructed from two first order ODEs which govern changes in immunogenic and tolerogenic programmes respectively. Each ODE is composed of 3 terms, with the regulatory influences modelled using Hill functions to describe sigmoidal associations. The first term describes auto-amplification of each programme; the second term describes the cross inhibition between opposing programmes; the final term allows for programme decay at a constant rate.

To make a more parsimonious model we assumed that the parameters that characterise generic interactions are constant (i.e. *a,b,k*=1, n=4 and θ=0.5) in accordance with these parameters creating a stable attractor landscape containing 3 states as described in (Huang et al.).

Analysis and plotting of the ODE model was performed within MATLAB (Mathworks, Inc.). Trajectories were found using the ode45 solver and phase portraits were produced using the quiver command. TF expression values or z-scores representing expression of multiple TFs in each single cell were exported from Scanpy scRNA-seq analysis, scaled to fit phase portrait boundaries and then utilised as time 0 starting points from which trajectories were calculated and plotted. The total number of cells trajectories ending at each of the 3 attractors after simulation was quantified and then plotted as pie charts in GraphPad Prism 8 software for comparison.

### Data Sharing Statement

figgFor original data, please contact. RNA-sequencing data are available at GEO under accession number GSE166079.

## Acknowledgments

We acknowledge the use of the IRIDIS High Performance Computing Facility and Flow Cytometry Core Facilities, together with support services at the University of Southampton. The study was funded by a Sir Hendy Dale Fellowship from Wellcome Trust, 109377/Z/15/Z. Development of single cell Drop-Seq technology was funded by MRC grant MC_PC_15078. Foreskin LC isolation was funded by a South African NRF Thuthuka funding grant, TTK150624120787.

---

**Supplementary Table 1**. Gene lists of the LC immunogenic signature derived from (Sirvent et al. 2020) and tolerogenic signature derived from (Davies et al. 2019).

**Supplementary Table 2**. Toggle-switch model iterations utilising different combinations of immunogenic vs tolerogenic defining TFs in LCs.

**Supplementary figure 1.**
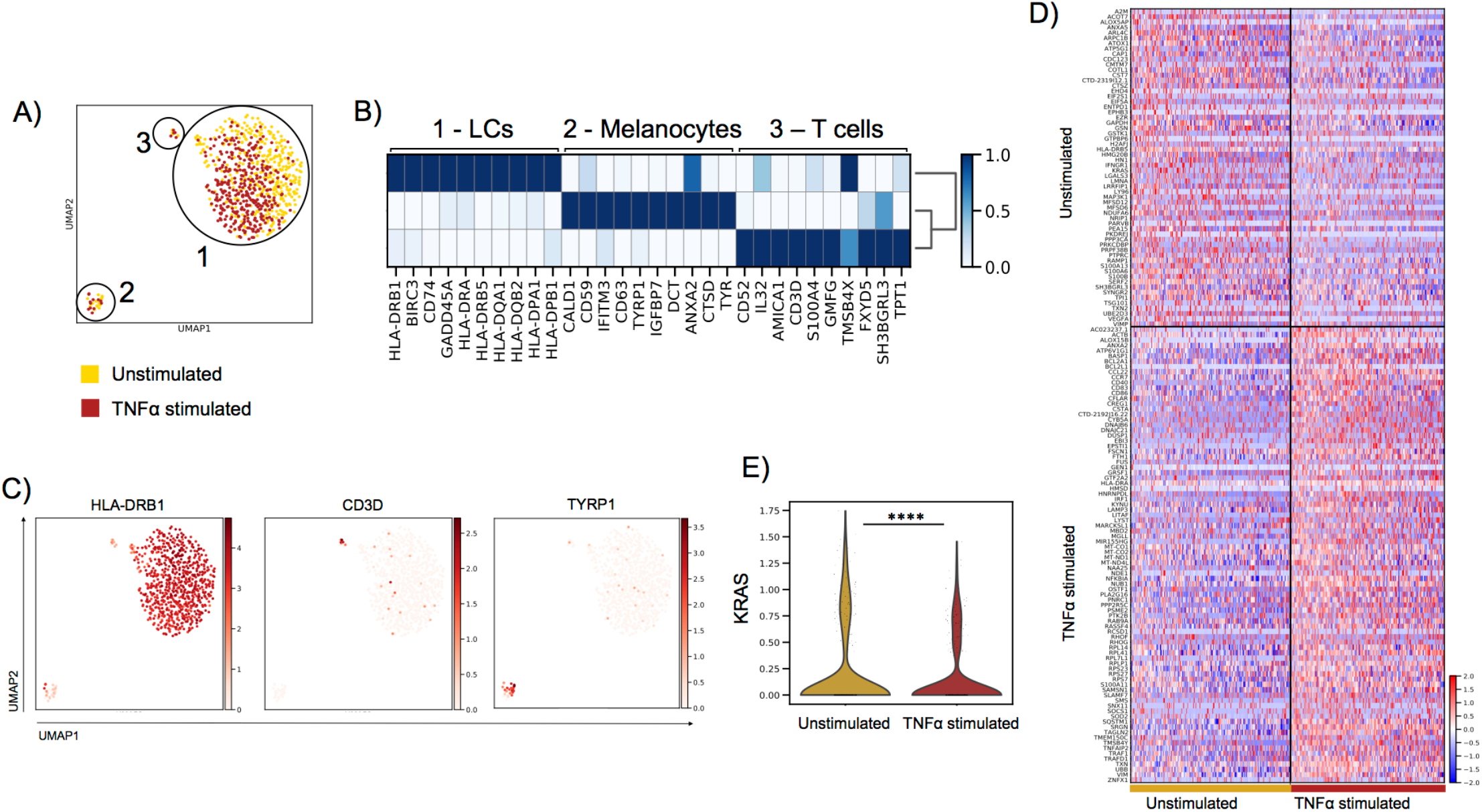
TNF enhances immunogenic transcriptional programming in migratory LC. **A**. UMAP dimensionality reduction analysis of epidermal cell populations detected 3 distinct subclusters of cells. **B**. Top 10 markers genes for clusters 1-3 (t-test, ScanPy pipeline, version=1.5.0), revealed populations to be LCs (cluster 1), melanocytes (cluster 2) and T cells (cluster 3) **C**. UMAP marker plots displaying the expression of the LC marker *HLA-DRB1*, the T cell marker *CD3D* and the melanocyte marker *TYRP1*. **D**. Heatmap displaying the 61 upregulated DEGs in unstimulated migrated LCs and 87 upregulated DEGs in TNF stimulated migrated LCs (FDR corrected p=<0.01, logFC>1). Gene ontology analysis (Toppgene) results are displayed alongside for unstimulated and TNF stimulated migrated LC upregulated DEGs (-log10 FDR corrected p-values) **E**. Violin plot of *KRAS* expression in unstimulated and TNFα stimulated migrated LC.

**Supplementary figure 2.**
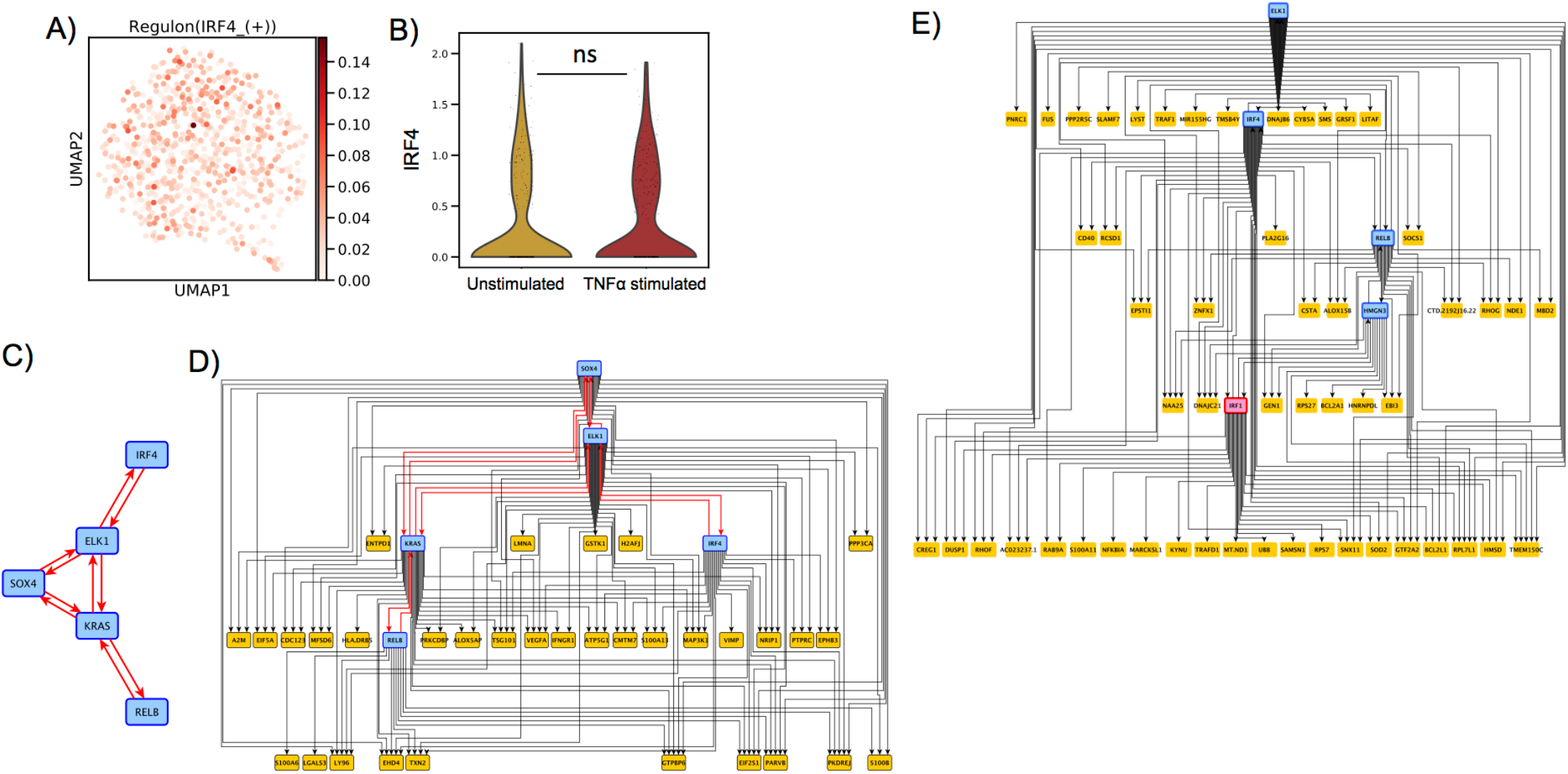
IRF1 expression controls immunogenic transcriptional programming. **A**. UMAP marker plot displaying *IRF4* regulon enhancement (z-scores) within individual LCs. **B**. Violin plot of *IRF4* expression in unstimulated and TNF stimulated migrated LCs. MAST, ****=p<0.001. **C**. PIDC network graph displaying connectivity (edge weight >1.5) between a regulatory module comprising of *SOX4, KRAS, IRF4, RELB* and *ELK1* in unstimulated migrated LCs. **D**. PIDC network graph comprising 38 nodes (5 TFs, 33 output genes) and 107 edges with weight >1.5, hierarchically organized, displaying predicted regulatory modules for the regulatory TF module from Supplementary Figure 2C along with upregulated DEGs in unstimulated LCs. **E**. PIDC network graph comprising 58 nodes (5 TFs, **C**. 53 output genes) and 122 edges with weight >1.5, hierarchically organized, displaying predicted regulatory modules for *IRF1* and TFs core to migrated LC (*IRF4, HMGN3, ELK1* and *RELB*), along with upregulated DEGs in TNF stimulated migrated LCs.

